# Serotonin attenuates tumor-necrosis-factor-alpha-induced intestinal inflammation by interacting with human mucosal tissue

**DOI:** 10.1101/2024.05.05.592559

**Authors:** Veronika Bosáková, Ioanna Papatheodorou, Filip Kafka, Zuzana Tomášiková, Marcela Hortová Kohoutková, Jan Frič

## Abstract

The intestine houses the largest reservoir of immune cells and is serviced by the largest and most complex peripheral nervous system in the human body. The gut-brain axis orchestrates bidirectional communication between the central and enteric nervous systems, playing a pivotal role in regulating overall body function and intestinal homeostasis. Using a human 3D *in vitro* model, we investigated the effect of serotonin, a neuromodulator produced in the gut, on immune cell and intestinal tissue interactions. Our findings revealed that serotonin attenuates the tumor-necrosis-factor-alpha-induced pro-inflammatory response, mostly by affecting the expression of chemokines. Serotonin was found to impact tissue-migrating monocytes’ phenotype and distribution, without direct contact with the cells, by remodeling the intestinal tissue. Collectively, using fully human 3D model of intestine, our results show for the first time that serotonin has a crucial role in communication among gut-brain axis components and regulates monocyte migration and plasticity, thereby contributing to gut homeostasis and the progression of intestinal inflammation. *In vivo* studies focused on role of neuromodulators in gut homeostasis and inflammation have shown controversial results, highlighting importance of development of human experimental models. Moreover, our results emphasize importance of human health research in human-cell-based models and suggests serotonin signaling pathway as new potential therapeutic target for inflammatory bowel disease patients.

## INTRODUCTION

The gut-brain axis is defined as comprising bidirectional interactions between the central nervous system and the largest and most complex component of the peripheral nervous system—the enteric nervous system (ENS). As the gut mucosa is able to respond to neural signaling (1), and intestinal enteroendocrine cells (EECs) represent a major source of some neuromodulators (2–7), interactions between intestinal tissue and the ENS are key mechanisms of gut homeostasis maintenance. How interactions between the intestinal tissue and the gut microbiome and the dysregulation of the mucosal immune system lead to an uncontrolled inflammatory response has been studied; however, the involvement of the ENS in these processes is not well described.

Intestinal tissue harbors the largest reservoir of immune cells in the human body (8); however, the interconnectivity of neuromodulators and the intestinal immune system has not been completely elucidated. Macrophages (Mϕ) are abundant in the intestinal tissue and exhibit remarkable functional plasticity in response to their microenvironment. Mϕ contribute to tissue homeostasis maintenance but can exacerbate inflammation when dysregulated (9). Intestinal Mϕ mainly originate from embryo-derived precursors and have a self-maintenance ability (9–12). However, the mature Mϕ pool can be also replenished by circulating monocytes that migrate into mucosal tissue (9,13). Under a pathological state, such as inflammatory bowel disease (IBD), monocyte-Mϕ differentiation is dysregulated, leading to impaired bacterial clearance and the production of cytokines such as tumor necrosis factor (TNF)α (14–18), exacerbating tissue damage. Therapeutic inhibition of TNFα has revolutionized management of IBD, proving that TNFα has a crucial role in pathogenesis of intestinal inflammation. The role of TNFα has been well studied in the context of inflammation, which it promotes by inducing cell death (19). However, the effect of TNFα stimulation on mucosal tissue and its connection to the gut-brain axis is still not fully understood because of the lack of suitable immune-cell-depleted tissue and ENS models. Moreover, the development of MΦ differs in mice and humans, and studies are impeded by the difficulty in creating human cell models. Therefore, how mucosal tissue influences monocyte migration, fate, and differentiation in humans is still unclear.

Neuromodulators have crucial roles in the regulation of physiological functions and homeostasis in the human body, with the latter affecting multiple systems, including the nervous system, mucosal tissue, and the immune system. Various cells, including enteric neurons, immune cells, gut microflora, and intestinal EECs, are sources of neuromodulators. In particular, 50% of whole-body dopamine and more than 95% of total serotonin are produced in the gut, mainly by EECs (2–7). As well as their well-established roles in the nervous system, dopamine and serotonin exert pleiotropic effects in intestinal tissues (2), and both are involved in the inflammation and pathogenesis of IBD (2,20). The roles of serotonin in the development and progression of intestinal inflammation and related pathologies have been thoroughly studied and reviewed (20–23). A variety of immune cells, including monocytes, express serotonin receptors and thus respond to serotonin stimulation (24,25). In this context, serotonin stimulation is believed to promote monocyte chemotaxis towards sites of inflammation (26); however, the precise mechanisms mediating these actions are unresolved.

Animal models have been widely used to investigate the mechanisms and pathogenesis of human inflammatory diseases. Though such models have generated important findings, rodents and humans differ in their susceptibility to pathogens and microbiome composition. In mucosal tissue, TLR signaling is a vital part of innate immunity. Furthermore, given the ligand recognition-specificity of each TLR, the expression and localization of the receptors are crucial for an adequate immune response. However, the expression of TLRs differs strikingly between mice and humans throughout the whole gastrointestinal tract (27), and therefore the translation of *in vivo* results can be challenging. In context of serotonin, controversial findings were found when mouse and rat models were used to elucidate role of this neuromodulator in intestinal inflammation (28–30). Therefore, human-cell-based intestinal organoid model provides a solution in order to elucidate these conflicting results obtained with animal models. Organoids are 3D “organ in a dish” models that aim to recreate key aspects of the *in vivo* structure of tissues using a multiple cell types from the species of interest, generating a simulated microenvironment in which the organoid cells exhibit key aspects of organ function (31). Previously, we have shown that induced pluripotent stem cell (iPSC)-derived intestinal organoids (IOs) form an immunocompetent environment, facilitating the co-cultivation of immune cells and observations of their interactions with mucosal tissue (32). Therefore, IOs represent potent models for studying the roles of neuromodulators in homeostasis and the immune functions of intestinal mucosal tissue.

Despite growing evidence highlighting the crucial role of neuromodulators, particularly serotonin, in intestinal pathophysiology, the underlying mechanisms of action are elusive. Here, we aimed to describe effect of TNFα stimulation on mucosal tissue in 3D human *in vitro* model. We hypothesized that serotonin is involved in development and resolution of intestinal inflammation affecting both mucosal tissue and resident immune cells. Therefore, we aimed to describe the role of serotonin in both homeostasis and TNFα-induced inflammation using human-cell-based model overcoming thus limitations and interspecies differences of *in vivo* models. Current therapeutic strategies for IBD rely on attenuating of intestinal inflammation using immunosuppressants, immunomodulators and biological treatment targeting specific cytokines or immune cell types. Despite the progress of development of new agents, significant portion of IBD patients show as primarily or secondary nonresponsive to conventional treatment. Therefore, investigation of novel therapeutic approaches is emerging.

In this study, we elucidated the impact of neuromodulators, including serotonin, on the expression profiles of human gut mucosa cells and infiltrating monocytes. We used state-of-the-art 3D *in vitro* organoid models depleted of all ENS and immune cells and the *ex vivo* co-cultivation of IOs with human monocytes. For the first time, the direct effects of serotonin on mucosal tissue during homeostasis and TNFα-induced inflammation were studied. Our study has shed light on the mechanisms underlying the role of serotonin in human gut pathophysiology in the context of inflammation and provided clues to the bidirectional communication occurring in the gut-brain axis. Altogether, our results suggest that therapeutic targeting serotonin might alleviate intestinal inflammation in IBD patients.

## RESULTS

### Organoids form a complex environment for studying mucosal homeostasis

Mucosal tissue consists of epithelial cells that abundantly express E-cadherin and form a polarized layer, with a brush border enriched in F-actin facing the lumen of the gut. In this study, we used human iPSC-derived IOs to simulate mucosal tissue. Firstly, to map the complex structures of the IOs, we used immunofluorescence (IF) labelling of F-actin (phalloidin) and E-cadherin (**Fig. 1A**). We investigated the polarization of the tissue using hematoxylin and eosin (**Fig. 1B**), and visualized the presence of organized structures, with the inner/lumen side of the organoid facing a lumen. Immunohistochemistry labeling showed the expression of MUC5AC by the epithelial cells (**Fig. 1C**), indicating the mucus-producing functionality of the IO tissue. Next, flow cytometry analysis was employed to determine the presence of epithelial (CD90^−^EpCam^+^) cells and mesenchymal (CD90^+^EpCam^−^) cells (**Fig. 1D left**), as well as the absence of immune cells, as demonstrated by the lack of CD45^+^ cells (**Fig. 1D right**). Using IF labelling, we further verified that the epithelial and mesenchymal cells had formed organized structures within the IOs (**Fig. 1E**), allowing us to study the homeostasis of the epithelial barrier. According to the results, the IOs contained cells expressing chromogranin A (CGA), a marker of EECs (**Fig. 1F**), and lysozyme, a marker of Paneth cells (**Fig. 1G**), demonstrating each IO’s relevance as a model to study the gut-brain axis. Altogether, the results demonstrate that the IOs resembled *in vivo* human intestinal tissue in their structure and functionality, facilitating our study of the mechanisms of homeostasis maintenance and communication within the gut-brain axis in healthy mucosal tissue.

**Fig. 1:**
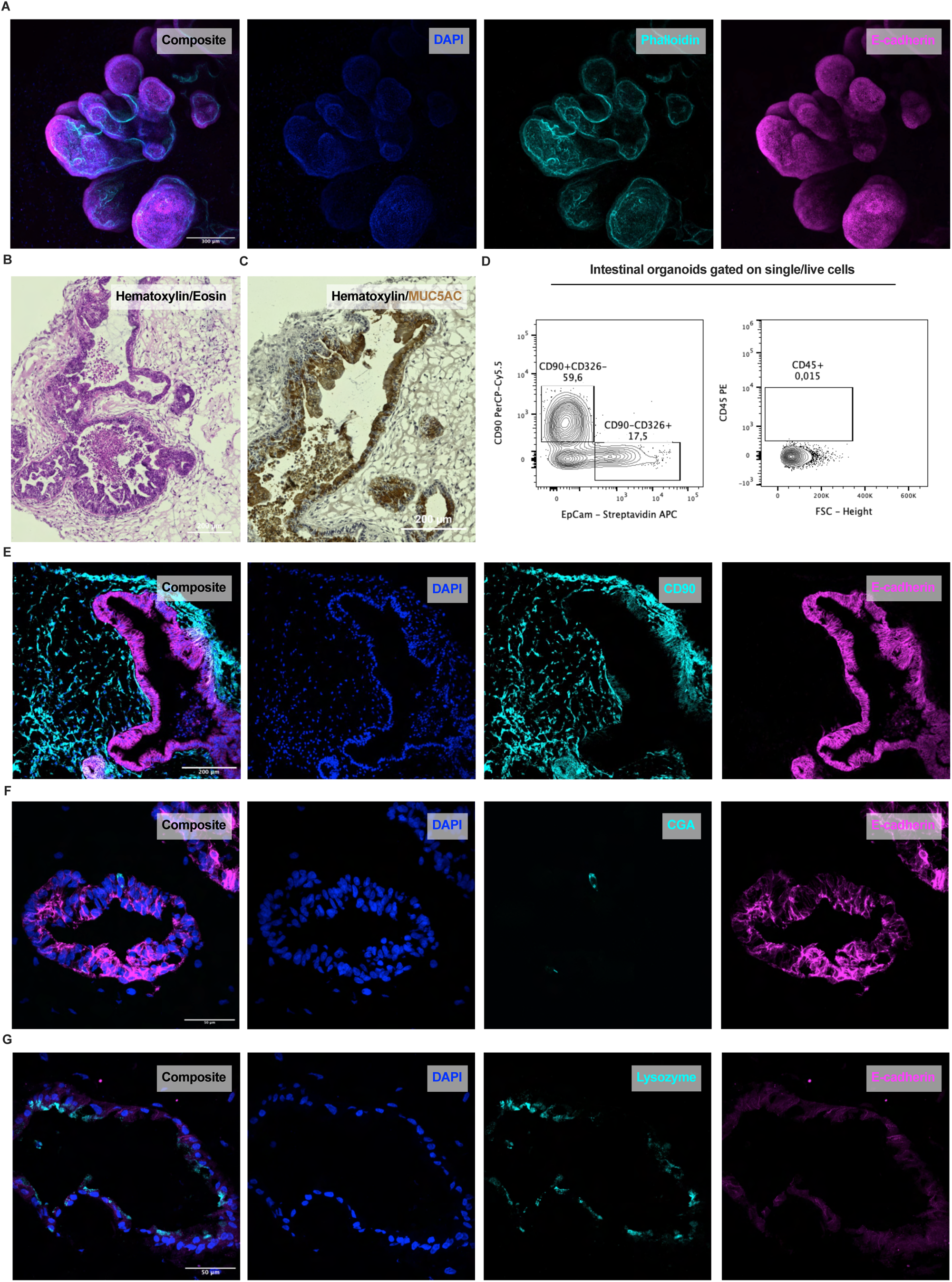
iPSC-derived organoids form complex organized structures containing enteroendocrine and Paneth cells. **A** Whole-mount immunofluorescence labelling for E-cadherin (magenta) and phalloidin (cyan), counterstained with DAPI (blue) depicting the polarity of IOs, scale bar 300 µm. **B** Hematoxylin and eosin staining demonstrating the structural organization of IOs, scale bar 200 µm. **C** Immunohistochemical labelling of MUC5AC counterstained with hematoxylin, scale bar 200 µm. **D** Flow cytometry analysis revealing expression of CD90, EpCam, and CD45 in IO cells. **E-G** Immunofluorescence labelling of E-cadherin (magenta), scale bar 200 µm, CD90 (cyan), CGA (cyan), scale bar 50 µm, and lysozyme (cyan), scale bar 50 µm, counterstained with DAPI (blue), showing epithelial, mesenchymal, enteroendocrine, and Paneth cells within IOs.

### TNFα stimulation of intestinal organoids initiates chemotaxis and pro-inflammatory cytokine response in intestinal tissue

Next, TNFα, one of the main drivers of inflammatory responses, was used to stimulate the IOs to define their capacity to model the progression of inflammation in mucosal tissue. IOs were treated with human recombinant (hr)TNFα for 4 hours and, subsequently, global transcriptomic alterations were assessed by bulk RNA sequencing (RNAseq) (**Fig. 2A**). Differential expression analysis of the RNAseq data revealed 1,415 differentially expressed genes (DEGs), of which 496 (35%) were downregulated and 919 (65%) were upregulated in the TNFα-treated IOs compared to the untreated controls (**Fig. 2B, C**). Gene ontology (GO) analysis identified several enriched pathways relevant to the response of IOs to TNFα stimulation, including TNFα-mediated signaling and I-κB kinase/NF-κB signaling pathways (**Fig. 2D**). Interestingly, pathways associated with the chemotaxis of immune cells, such as leukocyte chemotaxis, monocyte chemotaxis, neutrophil chemotaxis, and chemokine production (**Fig. 2D**), were also enriched in the IOs treated with TNFα, with the most abundant DEGs in these pathways being CCL2, CCL11, CCL20, CXCL2, and CXCL8 (**Fig. 2E**). These results illustrate the important role of mucosal cells in shaping the progression of inflammation, which they achieve via modulation of immune cell chemoattraction signals. Toll-like receptor (TLR) signaling is an important initiator of the inflammatory response, inducing the expression of pro-inflammatory cytokines and chemokines to attract immune cells to the site of inflammation (33). Thus, we sought to investigate the expression of TLRs in the IOs. We observed high expression of TLR2, TLR3, TLR4, TLR5, and TLR6 (**Fig. 2F**), as expected for human intestinal tissue. The above findings demonstrate that the IOs provided relevant models to facilitate our study of intestinal inflammation and the importance of TLR signaling and subsequent alterations in chemotaxis in mucosal cell inflammatory responses. Thus, we hypothesized that TLR signaling and downstream processes in the mucosal layer might be controlled by neuromodulators.

**Fig. 2:**
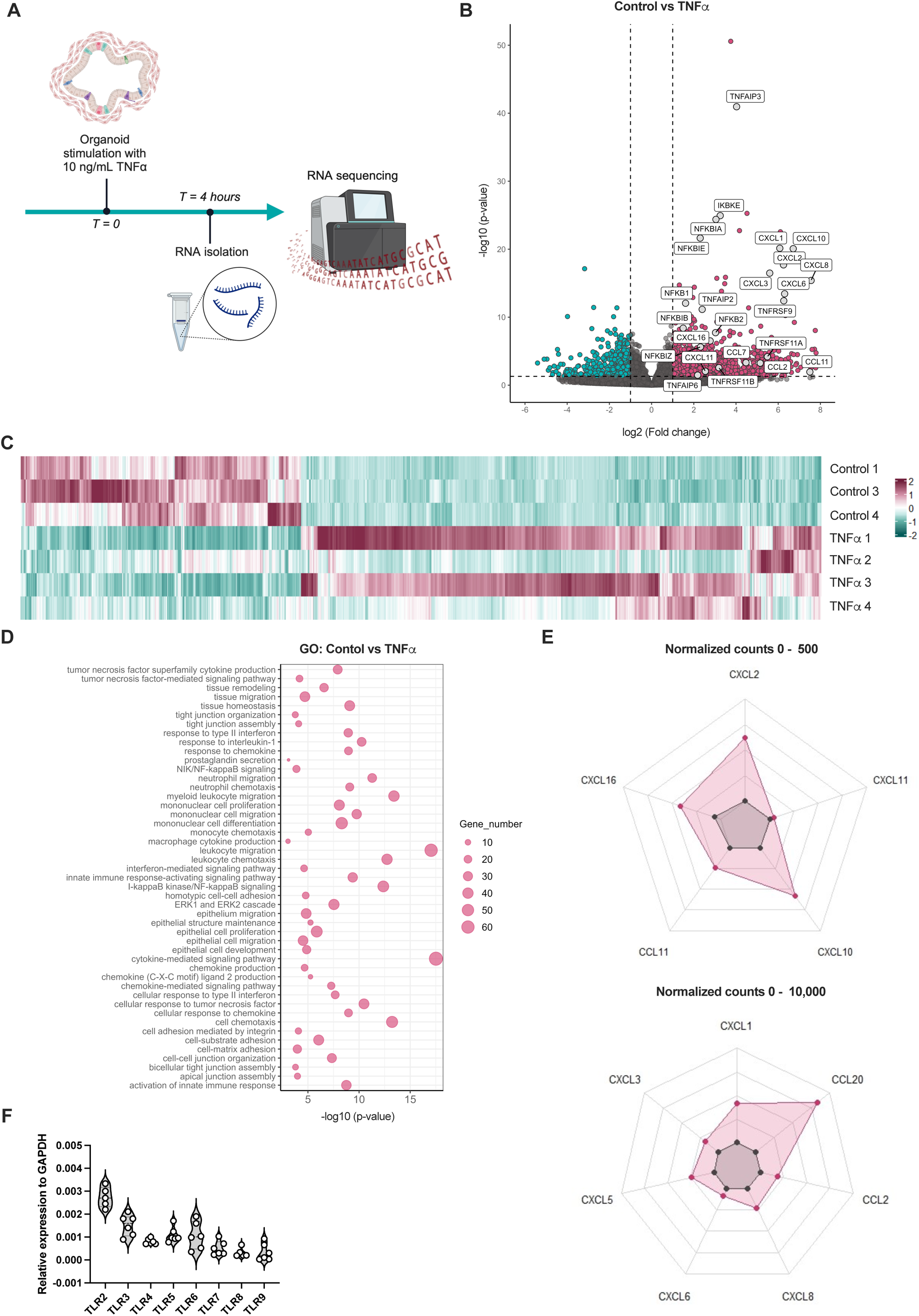
TNFα induces a pro-inflammatory environment in organoid tissue. **A** Scheme of *in vitro* experiments where IOS were stimulated with TNFα for 4 hours and expressional changes were analyzed using RNAseq. **B** Volcano plot depicting DEGs in TNFα-treated samples compared to untreated control. Magenta and cyan dots represent DEGs with *p* < 0.05. Vertical dashed lines represent the |log2 (Fold change)| = 1, and the horizontal dashed line *p*-value = 0.05. **C** Heatmap showing scaled normalized counts of DEGs of control and TNFα-treated samples. **D** Bubble chart demonstrating the most significantly enriched pathways obtained using gene ontology. **E** Spider plots showing the mean of normalized counts, between 0 and 500 and 0 and 10,000, of chemokine genes among significant DEGs. **F** Violin plot depicting Toll-like receptor RNA expression in IOs, dashed lines represent the median and dotted lines represent quartiles, N=6.

### Neuromodulators regulate Toll-like receptor expression in intestinal organoids

On consideration of the findings so far, we sought to determine the potential effects of neuromodulators on TLR signaling and the subsequent pro-inflammatory processes in mucosal tissue. Firstly, with our RNAseq data, we ascertained that the IOs expressed the neuromodulator receptors ADRB2, CHRNA1, CHRNB1, CHRNA5, CHRNA10, and HTRA2 (**Fig. 3A**). Next, we sought to determine if relevant neuromodulators alter TLR signaling or stimulate the inflammatory response in mucosal IOs under physiological conditions. For this purpose, we stimulated the organoids with the neuromodulators serotonin, dopamine, noradrenaline, and acetylcholine. Firstly, we used an RT^2^ Profiler PCR Array (Human Inflammatory Response & Autoimmunity) for screening the responses to the mentioned neuromodulators (**Fig. 3B**). Interestingly, the results indicated that the expression of various TLRs was decreased in presence of different neuromodulators. We corroborated this finding by determining the RNA expression levels of the TLRs in IOs untreated or treated with serotonin, dopamine, noradrenaline, or acetylcholine using quantitative (q)PCR. After stimulation with serotonin, the IOs showed significantly decreased expression of TLR2, TLR3, and TLR7 (**Fig. 3C**), no change in the expression of TLR5, TLR8 and TLR9 (**Fig. 3D**), and trend of decreased TLR4 and TLR6 (**Fig. 3E**) expression. We also observed a significant decrease in TLR7 (**Fig. 3C**) expression after acetylcholine stimulation. We confirmed the above findings at the protein level using IF staining of histological slides of IOs (**Fig. 3F-J**), focusing on TLR2, TLR3, TLR4, and TLR6, as the most highly expressed TLRs in IOs (**Fig. 2F**). Quantification of the IF revealed a significant decrease in TLR4 and TLR6 expression in all treatment groups compared to untreated controls (**Fig. 3F**). We also observed a significant decrease in TLR2 (**Fig. 3G**) and a trend of decreased TLR3 expression after serotonin and dopamine treatment (**Fig. 3H**). Given the abundance of TLR2, TLR3, TLR4, and TLR6, as shown in **Fig. 2F**, we concluded that neuromodulators governed important immune response pathways in the IOs, and thus *in vivo* mucosal tissue, by modulating the expression of TLRs.

**Fig. 3:**
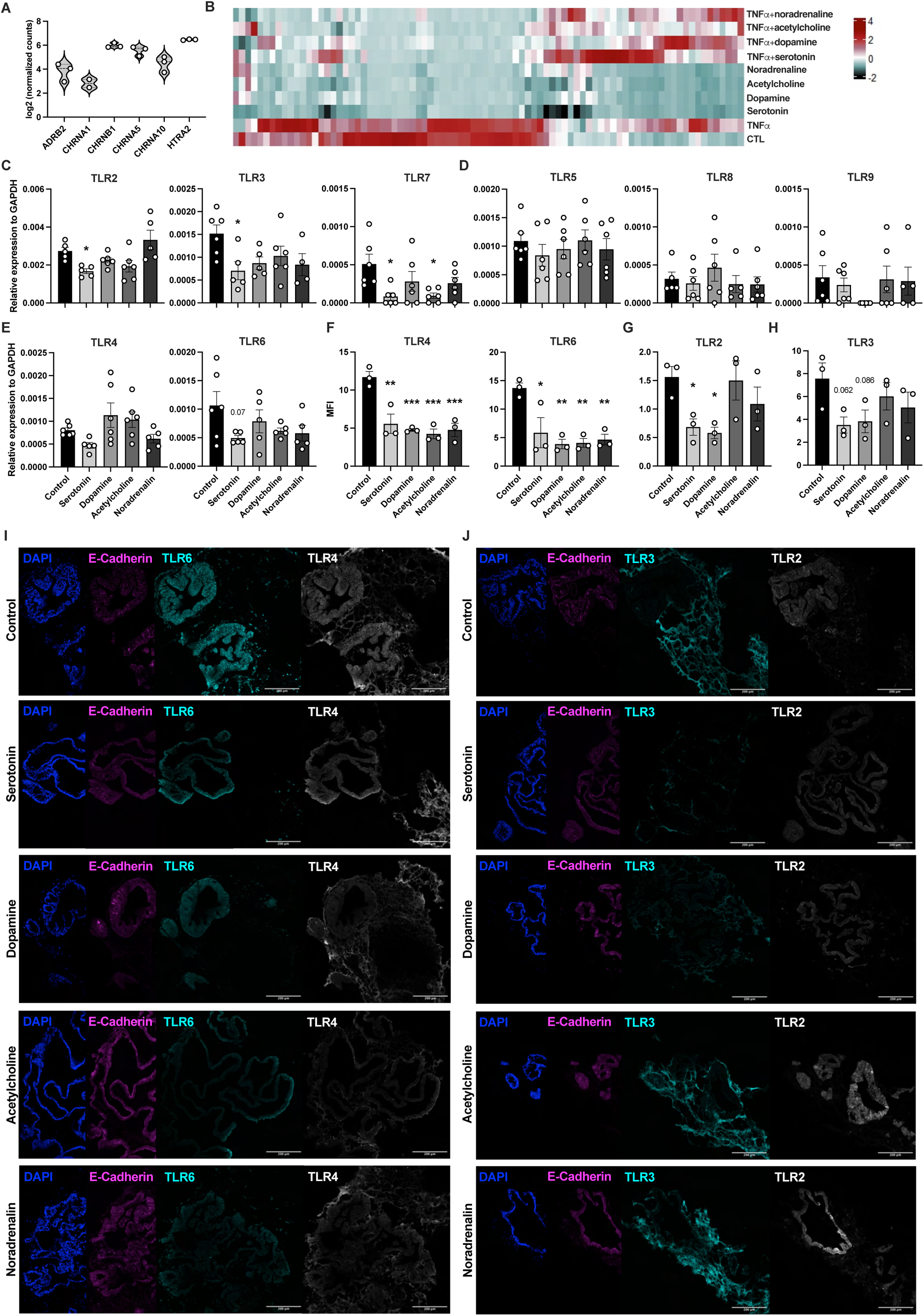
Neuromodulators regulate Toll-like receptor signaling in intestinal tissue. **A** Violin plot depicting RNA expression of neuromodulator receptors by IOs. Dashed lines represent the median and dotted lines represent quartiles, N=5-6. **B** Heatmap showing results from RT^2^ Profiler PCR Array (Human Inflammatory Response & Autoimmunity) gene expression screening for IOs after serotonin, dopamine, acetylcholine, and noradrenaline stimulation. **C-E** RNA expression of TLR2, TLR3, TLR6, and TLR7 of IOs treated with the indicated neuromodulators. Data are shown as mean ± SEM, N=4-6. **F-H** Quantification of mean fluorescence intensity of immunofluorescent labelling for TLR2, TLR3, TLR4, and TLR6 in IOs stimulated with neuromodulators. Data are shown as mean ± SEM, N=3. **I, J** Representative immunofluorescence labeling of TLR2, TLR3, TLR4, TLR6, and the epithelial cell marker E-cadherin in IOs untreated or treated with serotonin, dopamine, noradrenaline, or acetylcholine, counterstained with DAPI (blue), scale bar 200 µm, N=3. Statistical significance was determined by ordinary one-way ANOVA, \**p* < 0.05, \*\**p* < 0.01, \*\*\**p* < 0.005.

### Serotonin exerts a modest non-pro-inflammatory effect on healthy mucosal tissue

Next, we aimed to explore more deeply into the effects of neuromodulators on intestinal tissue homeostasis using our RNAseq dataset. As we observed serotonin to have the most prominent effect on the IOs, we decided to focus only on stimulation with serotonin. Firstly, to test the effect of serotonin on healthy mucosal tissue, we stimulated IOs with serotonin and compared them to untreated IOs as a control. Serotonin influenced healthy tissue by altering its gene expression pattern (**Fig. 4A**). However, the analysis revealed only 249 DEGs in our dataset, of which 211 (85%) were upregulated and 38 (15%) were downregulated after serotonin treatment compared to untreated control (**Fig. 4A, B**), suggesting that serotonin did in fact affect the tissue under physiological conditions. GO analysis of these DEGs revealed the enriched pathways in serotonin-treated samples were related to tissue remodeling and included tight junction assembly and cell-cell junction assembly (**Fig. 4C**). Therefore, it appears that, in healthy intestinal tissue, serotonin had a subtle effect, inducing the remodeling of epithelial barrier integrity through tight junction assembly.

**Fig. 4:**
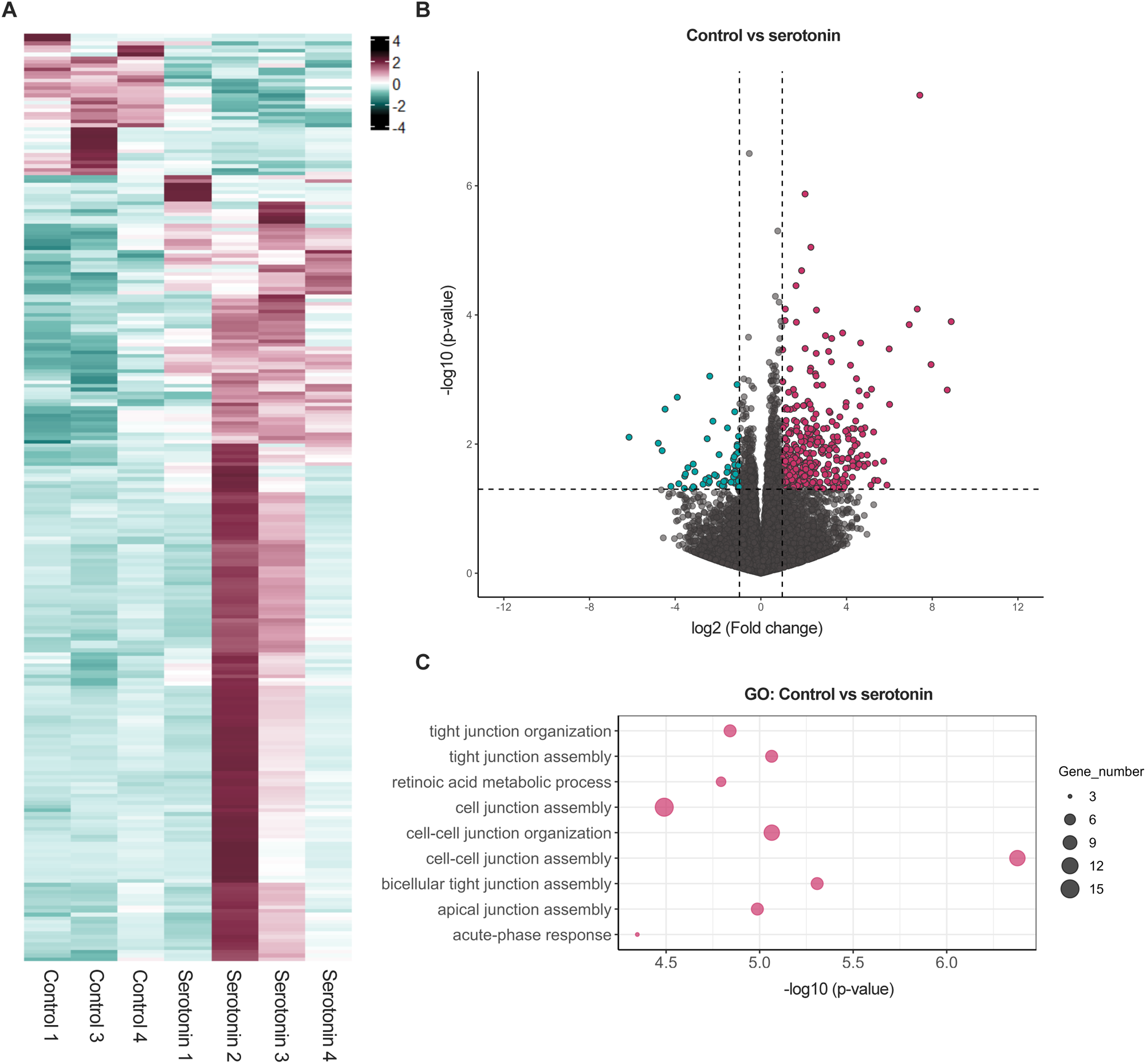
Serotonin has a mild effect on healthy intestinal tissue. **A** Heatmap showing scaled normalized counts of DEGs of control and serotonin-treated samples. **B** Volcano plot depicting DEGs in serotonin-treated IOs compared to untreated IOs. Magenta and cyan dots represent DEGs at *p* < 0.05. Vertical dashed lines represent the |log2 (Fold change)| = 1, and the horizontal dashed line *p*-value = 0.05. **C** Bubble chart showing the most significantly enriched pathways in serotonin-treated IOs compared to untreated controls obtained using gene ontology.

### Serotonin decreases chemokine expression in TNFα-induced intestinal inflammation

Thus far, our findings support the idea that serotonin exerts a modest effect on intestinal mucosal tissue under physiological conditions. We also showed that serotonin, along with other neuromodulators, affects the immune response by altering TLR expression in mucosal tissue. Interestingly, previous research has also shown the possible interplay between serotonin and TNFα-signaling pathways (30,34). Thus, we aimed to describe the role of serotonin in inflammation by taking advantage of both our IO model of TNFα-induced inflammation and our RNAseq dataset (**Fig. 5A**). Interestingly, in the presence of serotonin, we observed much fewer DEGs (695) than in the TNFα-only-treated IOs (1415). Furthermore, while the large majority of genes were upregulated (919 (65%)) in TNFα-treated organoids, in the presence of both TNFα and serotonin, most genes were downregulated (477 (69%)), suggesting that serotonin had an attenuating effect on TNFα stimulation. To confirm this, we determined the common and unique gene expression patterns between these two conditions (**Fig. 5B**). Strikingly, we observed that more than 30% of the genes upregulated in TNFα-treated samples were downregulated in the presence of serotonin, and more than 20% of the downregulated genes in TNFα-treated samples were upregulated in the presence of serotonin, reiterating that serotonin attenuated TNFα-induced inflammation. Next, we performed a detailed analysis of the dataset, comparing TNFα+serotonin-treated IOs and TNFα-treated IOs. The presence of serotonin induced a consistent response, as can be seen in the heatmap (**Fig. 5C**). Most of the DEGs in this comparison were downregulated (**Fig. 5D**). To reveal if the genes that were differentially abundant in the presence of serotonin participate in important pathways, we performed GO analysis. Again, the gene functions were mainly concentrated in pathways involved in the chemotaxis and migration of leukocytes (**Fig. 5E**), suggesting that serotonin potentially attenuates the course of TNFα-induced inflammation by influencing the movement of immune cells. Therefore, we focused on the main chemokines induced by TNFα and their expression changes in the presence of serotonin. We found a significant decrease in the expression of CCL2, CXCL1, CXCL5, CXCL6, and CXCL8 (**Fig. 5F**) in TNFα+serotonin-treated IOs compared with the TNFα-only group. Notably, none of these chemokines were affected by serotonin stimulation of healthy tissue (**Fig. 5F**).

**Fig. 5:**
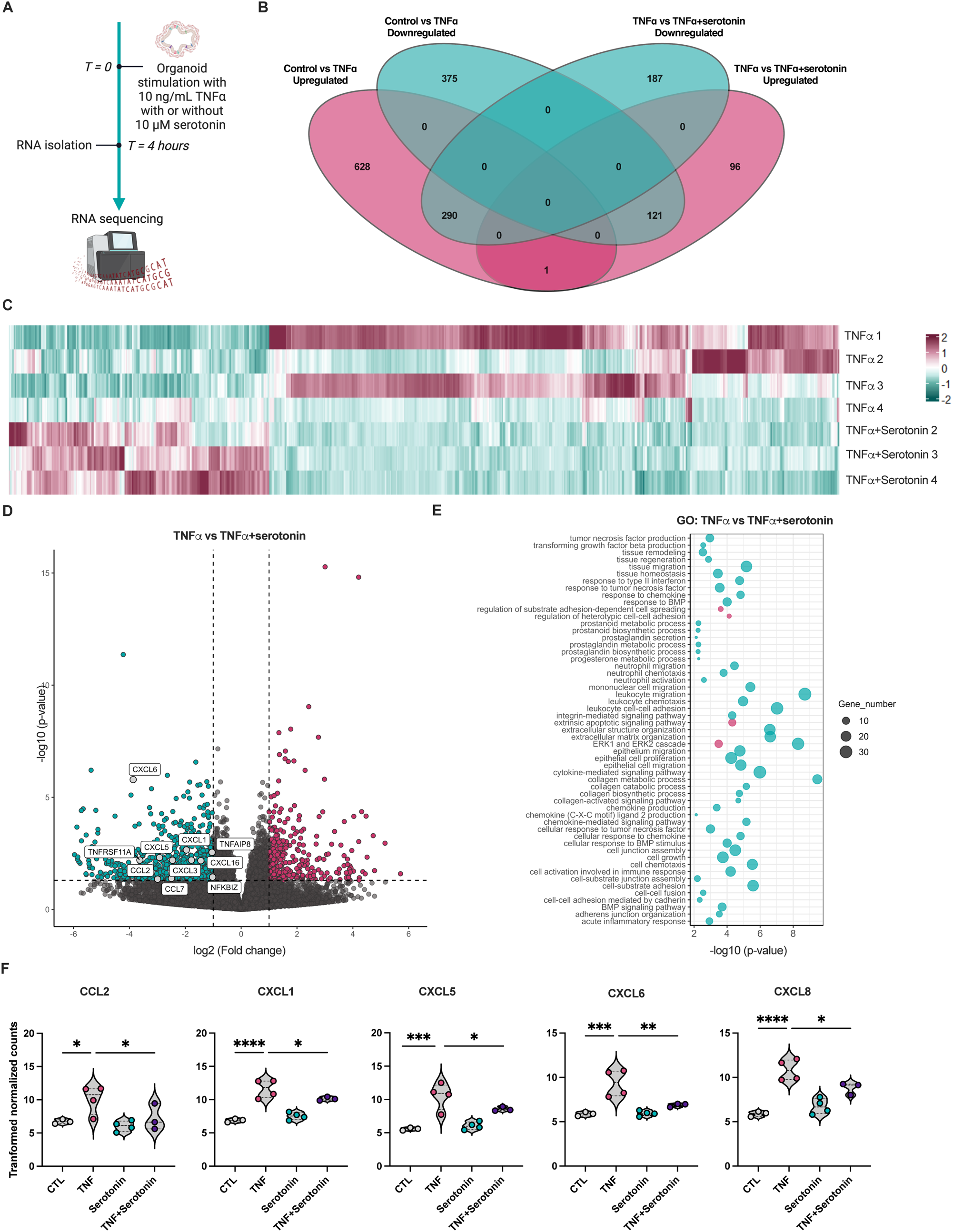
Serotonin affects the expression of chemokines in the pro-inflammatory state of intestinal tissue. **A** Schematic for *in vitro* experiments using IOs. IOs were stimulated with TNFα with or without serotonin for 4 hours, total expression changes were assessed using bulk RNAseq. **B** Venn diagram depicting DEGs in TNFα- and TNFα+serotonin-treated samples compared to TNFα-treated and untreated controls**. C** Heatmap showing scaled normalized counts of DEGs of control and serotonin-treated samples. **D** Volcano plot depicting DEGs in TNFα-treated IOs compared to TNFα+serotonin-treated IOs. Magenta and cyan dots represent DEGs with *p* < 0.05. Vertical dashed lines represent the |log2 (Fold change)| = 1, and the horizontal dashed line *p*-value = 0.05. **E** Bubble chart pinpointing the most significantly enriched pathways obtained using gene ontology. Magenta represents upregulated pathways and cyan represents downregulated pathways**. G** Violin plots showing expression of CCL2, CXCL1, CXCL5, CXCL6, and CXCL8 in control (CTL) and TNFα-, serotonin-, TNFα+serotonin-treated samples. Dashed lines represent the median and dotted lines represent quartiles, N=3-4. Statistical significance was determined by ordinary one-way ANOVA, \**p* < 0.05, \*\**p* < 0.01, \*\*\**p* < 0.005, \*\*\*\**p* < 0.001.

### Serotonin alters intestinal tissue’s ability to attract monocytes during TNFα-induced inflammation

To determine if the serotonin-induced changes in inflammatory cell pathway genes translate to functional effects, we performed chemotaxis and migration assays (**Fig. 6A**). To study how the chemokines released by IOs affect the migration of peripheral blood mononuclear cells (PBMCs), we seeded PBMCs from healthy donors into Transwell inserts and incubated them with IO supernatant treated with TNFα in the presence or absence of serotonin (**Fig. 6A upper part**). To determine the effects of serotonin on the chemoattraction of different subpopulations of immune cells, flow cytometry analysis was performed. The results indicated there were no differences in the chemoattraction of B cells (**Fig. 6B**), total T cells (**Fig. 6C**), CD4^+^ T cells (**Fig. 6D**), or CD8^+^ T cells (**Fig. 6E**) among the treatment groups. However, we observed the significantly decreased chemoattraction of classical (CD14^+^CD16^−^) (**Fig. 6F**) and increased chemoattraction of nonclassical (CD14^−^CD16^+^) (**Fig. 6G**) monocytes in the presence of supernatant from TNFα-treated IOs. The chemoattraction of intermediate (CD14^+^CD16^+^) monocytes was not affected by TNFα treatment (**Fig. 6H**). In agreement with our previous results (**Fig. 5F**), serotonin treatment of healthy tissue had no effect on the chemoattraction of any of the abovementioned cell types (**Fig. 6B-H**). However, when supernatant from IOs treated with both TNFα and serotonin was used, we observed a significant increase in classical (**Fig. 6F**) and a decrease in nonclassical (**Fig. 5G**) monocyte chemoattraction compared with the observations with TNFα-only supernatant. We also observed a significant decrease in the chemoattraction of intermediate monocytes when supernatant from TNFα+serotonin-treated samples was used compared to supernatant from serotonin-treated or untreated samples (**Fig. 5H**).

**Fig. 6:**
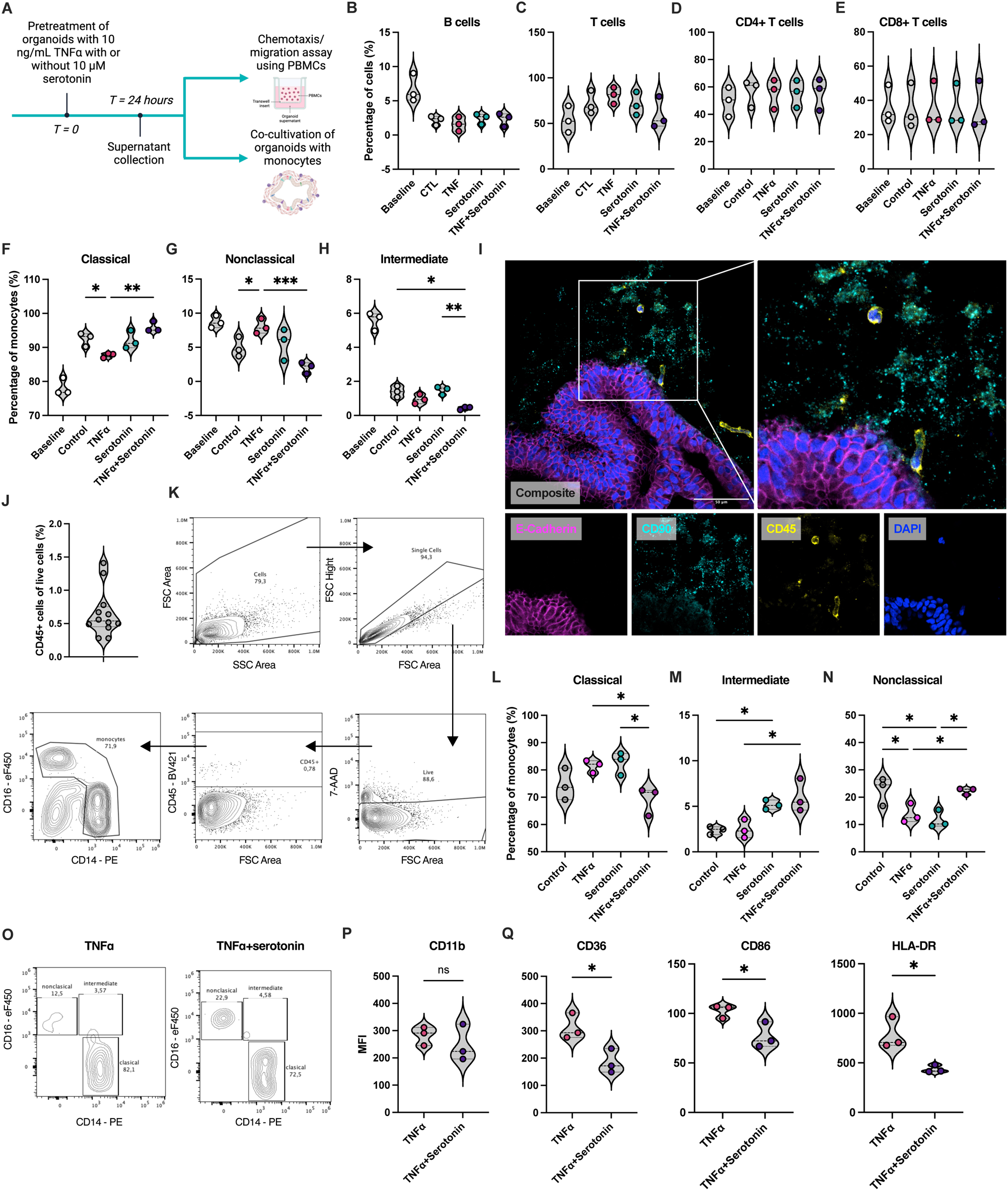
Serotonin influences the chemotaxis of monocytes and their fate in intestinal tissue in pro-inflammatory conditions. **A** Schematic for *in vitro* experiments focusing on chemotaxis and migration of PBMCs and monocytes. IOs were treated with TNFα in presence or absence serotonin, after 24 hours supernatants were collected and used for chemotaxis assay using Transwell inserts. Stimulated organoids were used for co-cultivation with monocytes. **B-E** Percentages of B cells, T cells, and CD4^+^, and CD8^+^ T cells that were chemoattracted and migrated towards supernatants of IOs untreated (CTL) or treated with TNFα, serotonin, or TNFα+serotonin. **F-H** Percentages of classical, non-classical, and intermediate monocytes that were chemoattracted and migrated towards supernatants of IOs untreated (control) or treated with TNFα, serotonin, or TNFα+serotonin, N=3. **I** Immunofluorescence labelling of E-cadherin, CD90, CD45 of IOs co-cultivated with monocytes, demonstrating their migration towards IO tissue, counterstained with DAPI (blue), scale bar 50 µm. **J** Percentage of CD45^+^ cells in live cells in flow cytometry analysis of dissociated IO cells co-cultivated with monocytes. Baseline represents percentages of respective cell types in total PBMCs labelled straight after isolation. **K** Representative contour plots of the gating strategy for monocytes co-cultivated with IOs. **L-M** Percentages of classical, non-classical, and intermediate monocytes cultivated with IOs stimulated with TNFα serotonin, or TNFα+serotonin, N=3. **O** Representative contour plots showing expression of CD16 and CD14 by monocytes co-cultivated with IOs pretreated with TNFα with or without serotonin. **P, Q** Expression of CD11b, CD36, CD86, and HLA-DR by monocytes co-cultivated with IOs stimulated with TNFα with or without serotonin, N=3. For violin plots, dashed lines represent the median and dotted lines represent quartiles. Statistical significance was determined by ordinary one-way ANOVA (B-E, F-H, L-N), and unpaired t-test (P, Q), \**p* < 0.05, \*\**p* < 0.01, \*\*\**p* < 0.005, ns, not significant.

### Serotonin remodels mucosal tissue to affect the phenotype and fate of tissue-migrating monocytes

Given the results on the effects of serotonin on the chemoattraction and migration of monocytes, the remodeling of healthy tissue, and the attenuation of inflammatory signaling pathways, such as TLR signaling, obtained in the previous steps, we sought to determine whether serotonin alters monocytic migration by remodeling IOs tissue. Our group has already demonstrated that IOs can be co-cultivated and interact with human monocytes (32). Therefore, we conducted the co-cultivation of monocytes isolated from healthy donors with IOs pretreated with TNFα with or without serotonin (**Fig. 6A lower part**). Firstly, we assessed the presence of monocytes migrating towards IO tissue with histological slides and IF staining. After 24 hours of co-cultivation, we were able to detect CD45^+^ cells within IO tissue in close proximity to mesenchymal and epithelial cells (**Fig. 6I**). Flow cytometry analysis of dissociated IOs co-cultivated with monocytes revealed the presence of CD45^+^ cells (0.6255 ± 0.3514%) (**Fig. 6J**), which were further identified as monocytes using CD14 and CD16 markers (**Fig. 6K**). When the percentages of monocytic subpopulations in the presence or absence of TNFα were determined, we observed no alterations in the migration of classical (**Fig. 6L**) or intermediate (**Fig. 6M**) monocytes and a decrease in the migration of nonclassical monocytes (**Fig. 6N**) towards IOs. Interestingly, we observed a significant decrease in classical (**Fig. 6L, O**) and an increase of intermediate and nonclassical (**Fig. 6M-O**) monocyte migration in TNFα+serotonin-pretreated compared to TNFα-pretreated IOs. We also observed an increase in intermediate (**Fig. 6M**) and a decrease in nonclassical (**Fig. 6N**) monocyte migration toward the intestinal tissue when the IOs were pretreated with serotonin alone. Considering the differences in the migration of the different populations of monocytes, we focused on the migratory and activation markers of the cells. Interestingly, we did not observe any significant differences in CD11b expression on monocytes when TNFα-pretreatment and TNFα+serotonin-pretreatment samples were compared (**Fig. 6P**). Conversely, we observed a significant decrease in CD36, CD86, and HLA-DR expression on monocytes that migrated towards IOs pretreated with TNFα+serotonin compared to those cultured with TNFα-only-treated IOs (**Fig. 6Q**). Overall, the evidence implies that serotonin affects healthy and inflamed intestinal tissue, resulting in alterations in the attraction of monocytes towards the tissue.

## DISCUSSION

The roles of neuromodulators in gut homeostasis and the different mechanisms involved have been comprehensively described (35). However, the mechanistic interconnections between neuromodulator signals in the human gut mucosa and the immune system are not as well known. As intestinal tissue is the main source of serotonin, we hypothesized that serotonin might affect immune response, migration of immune cells and the homeostasis in the mucosal tissue. Using state-of-the-art 3D, we aimed to create human-cell-based model of TNFα-induced inflammation and thus overcome limitation of animal and 2D cell line models. Here we aimed to study effect of neuromodulators, especially serotonin on mucosal tissue, resident immune cells, and its role in inflammation control and resolve the controversies in mouse and rat models showing its opposing effects.

Here, we showed that IOs possess distinct expression patterns of TLRs, with TLR2 and TLR6 being mostly expressed by epithelial cells, TLR3 mainly found in mesenchymal cells, and TLR4 expressed both in epithelial and mesenchymal cells. To this date, only the mouse intestine TLR expression map is available (36). Interestingly, compared to this map, we observed different TLR localization within the IOs, pinpointing inter-species differences between mouse and human intestinal tissue. The influence of microbiota on intestinal neural tissue and its production of neuromodulators has been intensively studied and reviewed (37–39). However, how neuromodulators affect expression of TLRs and thus the sensing of microbiota is not known. In this study, we showed that stimulation of IOs with neuromodulators, namely dopamine, noradrenaline, acetylcholine, and serotonin, induced a decrease in the expression of various TLRs, including TLR2 and TLR4. Interestingly, we observed different effects of neuromodulator stimulation on the mRNA and protein levels of TLR expression in IOs, suggesting the direct regulation of innate signaling by these neuromodulators. Our results also suggest that the decreased dopamine and serotonin levels observed in inflamed tissue in IBD patients (40,41) might be linked to an altered expression of TLRs in these patients. RNA and protein quantification in intestinal tissue from cohorts of both pediatric and adult IBD patients revealed significant increase of TLR2 and TLR4 expression in patient groups compared to healthy controls (42–44). However, further research is needed to shed more light on the effect of neuromodulators on the expression and signaling of TLRs in different cell types within human mucosal tissue.

Previously, we have shown that iPSC-derived organoids do not consist of any immune cells (32) and thus enable the study of specific mechanisms of action of pro-inflammatory cytokines in mucosal tissue, which is challenging with conventional *in vivo* and *ex vivo* models. We focused on TNFα and its effect on intestinal tissue using our organoid model. During inflammation, TNFα is mostly produced by immune cells, with intestinal-tissue-resident MΦ and tissue-migrating monocytes being the major sources (45). However, the response of intestinal tissue to TNFα and the subsequent effects on the immune response is still poorly understood. A pilot study, in which mouse jejunal organoids were stimulated with TNFα, demonstrated the induced expression of CXCL2 (MIP-2), emphasizing the important role of intestinal tissue in responses to pro-inflammatory stimuli (46). In accordance with this, our study demonstrated that, upon TNFα stimulation, mucosal tissue represents a crucial source of various chemokines and, therefore, plays a pivotal role in the control of the inflammatory response through the chemoattraction of immune cells. After TNFα treatment, there was an increase in the expression of other chemokines important for the chemotaxis of innate and adaptive immune cells, including CXCL8 (IL-8), CCL2 (MCP-1), CCL11 (eotaxin-1), and CCL20. A previous study showed that short-term stimulation with TNFα increased CXCL2 levels and decreased the mRNA expression of chromogranin A (CGA) (46), a neuroendocrine secretory protein produced by EECs. However, in our dataset we did not observe significant alteration of CGA after stimulation with TNFα, highlighting time-dependent effect of TNFα and more complex interaction with EECs. EECs produce a huge variety of hormones and neuromodulators, including serotonin (47), and are involved in the chronic inflammation of the intestine (48,49) and neurodegenerative disorders (37), suggesting the close interconnectivity between TNFα and serotonin in intestinal inflammation.

Previous studies have suggested a pro-inflammatory role for serotonin from evidence obtained in mouse models of intestinal inflammation where in IL-10 knock-out model depletion of the serotonin transporter (SERT) significantly worsen general health and intestinal inflammation (28). On the other, in dextran sulfate sodium (DSS)-induced colitis mouse model, reduction of serotonin by depletion of tryptophan hydroxylase 1 (THP1) gene decreased the severity of the colitis associated with decline of MΦ infiltration and significantly lower concentration of pro-inflammatory cytokine such as TNFα (29). In another *in vivo* study, depletion of SERT in mice was shown to affect the size of intestinal villi and enterocyte cell division, demonstrating the importance of serotonin as a physiologic regulator of intestinal growth under homeostatic conditions (50). Contrary to reports based on mouse models, our results indicate that serotonin does not induce a pro-inflammatory response in human mucosal tissue. In an *in vitro* rat model, the activation of one of the serotonin receptors on smooth muscle cells suppressed multiple responses to TNFα stimulation (30,34). The authors showed that an agonist of the 5-HT2A receptor attenuated the TNFα-induced pro-inflammatory response by decreasing the expression of IL-6, ICAM–1, and VCAM-1 (30). In our study, we observed similar findings in the *in vitro* human model, and our results demonstrated the distinct effect of serotonin on TNFα-induced inflammation, revealing that serotonin and TNFα signaling synergize to interfere with human gut pathophysiology. Our findings suggest that results obtained in mouse models need to be interpreted cautiously when translating to human health research. Importantly, this study implicate that rat models are probably more accurate as *in vivo* models in this context. Our study focused on the clinical relevance and effects of serotonin on healthy and inflamed intestinal tissue, and the findings indicate further research focusing on the molecular pathways involved in the interactions between serotonin and TNFα signaling is needed.

The exaggerated infiltration of the immune cells in gut tissue drives many pathologies and even exacerbates existing inflammation. Monocytes, in particular, are pluripotent plastic cells that can differentiate and perform various effector functions after migration to target tissues (12), thus influencing the course and resolution of inflammation in various human pathologies. Recent study using single-cell spatial transcriptomic showed presence of inflammation-dependent alternative (IDA) MΦ specifically in colon of IBD patients (51). IDA MΦ expressed neuregulin 1, molecule involved in epithelial cell expansion and differentiation, suggesting their role in epithelium regeneration during inflammation. Interestingly, authors showed that IDA MΦ were transcriptionally similar to MΦ derived from monocytes stimulated with M-CSF and serotonin (51). These results emphasize crucial role of tissue microenvironment on monocyte/MΦ differentiation. Our study has provided the first demonstration that serotonin can alter mucosal tissue pathology by favoring the migration of intermediate and nonclassical monocytes during TNFα-induced inflammation, without affecting the chemotactic ability of monocytes in healthy tissue. In particular, serotonin seems to upregulate the expression of genes involved in pathways such as cell-cell adhesion. Contrastingly, in TNFα-induced inflammation, serotonin exerts a pleiotropic effect on monocyte migration by altering both tissue chemokine expression and homeostasis to favor the migration of nonclassical monocytes. Compared to classical monocytes, nonclassical monocytes have distinct transcriptomic profiles and functions (52). In addition to their antigen-processing capabilities, they differ from classical monocytes by their metabolic profiles and association with wound-healing processes (53). Interestingly, the expression of CD11b, a molecule important for the adhesion and migration of monocytes and MΦ (54), was not affected in monocytes that migrated towards TNFα+serotonin-treated organoids. This suggests that the altered migration of distinct monocyte subsets is caused by different mechanisms. Contrarily, we observed a decrease in CD36, CD86, and HLA-DR expression on monocytes migrating towards tissue stimulated with TNFα in the presence of serotonin. CD36 is a scavenger receptor involved in many immune cell functions, such as phagocytosis and monocyte activation (55), while CD86 is a co-stimulatory molecule important for the proper activation of lymphocytes and thus plays a critical role in the adaptive immune response (56–58). Overall, our findings imply that serotonin can alter the profiles of immune cells and their migratory potential, even without direct contact with the cells. This in turn indicates that serotonin influences the interplay between intestinal tissue and both the innate and adaptive immune systems to regulate inflammation.

Here, we describe the first evidence for the impact of serotonin on human gut tissue, obtained using human 3D iPSC-derived IOs. Collectively, our findings have demonstrated a new role for serotonin as a mitigating agent of TNFα-induced inflammation in human gut-like tissue and provided novel mechanistic insights into both its direct effect on the gut and its indirect effect on the immune system when it comes into contact with gut tissue. We showed that serotonin modulates mucosal tissue to alter the migratory potential of monocytes and change the phenotype of the migrating cells. Therefore, it seems that serotonin is the link connecting the nervous system to intestinal tissue and monocytic/MΦ plasticity. Thus, it contributes not only to the bidirectional gut-brain axis but also constitutes a signal involved in three-way crosstalk between the gut, the immune system, and the brain. Nevertheless, further research is needed to uncover the molecular mechanisms behind the serotonin effects observed in our study with human-cell-based model. These findings can shed light on serotonin signaling pathway as new potential therapeutic target for IBD patients. Despite the advances in management of IBD, significant part of patients remains nonresponsive to the treatment, and therefore development of novel therapeutic strategies is still needed. SERT inhibitors are widely used as antidepressant drugs to treat anxiety and depression in IBD patients, furthermore evidence on their effect on IBD-related complications is emerging (59). Intriguingly, recent study showed that use of antidepressant including serotonin agonists and SERT inhibitors was associated with reduced risk of surgery in IBD patients (60). Our results highlight critical differences in mouse, rat, and human physiology, and we showed that human iPSC-derived organoids serve as a relevant model for future translation research focused on the effect of serotonin agonists and SERT inhibitors in alleviation of intestinal inflammation in IBD patients.

## Acknowledgments

We thank the technical support team of the Center for Translational Medicine for technical assistance. We acknowledge the CF Genomics CEITEC MU supported by the NCMG research infrastructure (LM2023067 funded by MEYS CR) for their support with obtaining scientific data presented in this paper. We would like to thank Dr. Jessica Tamanini from Insight Editing London for critical review and editing of the manuscript prior to submission. Graphical images were generated using BioRender.

## Author contributions

Conceptualization: VB, MHK, JF; Methodology: VB, IP, MHK, JF; Investigation: VB, FK, IP, ZT; Visualization: VB, IP; Funding acquisition: JF; Supervision: MHK, JF; Writing - original draft: VB; Writing - review and editing: all.

## MATERIALS AND METHODS

### Human induced pluripotent stem cell maintenance

Human induced pluripotent stem cells (hiPSCs) (WiCell, DF19-9-7T (61)) were cultured in Matrigel-coated tissue culture dishes and maintained in mTeSR Plus medium (StemCell Technologies) in incubator at 37 °C, 5% CO_2_ and 95% humidity. The medium was changed every second day. When the cells reached ∼80% confluence, they were passaged using TryPLE (Gibco), and exposed to the RHO/ROCK pathway inhibitor (10 μM, StemCell Technologies) for 24 hours in incubator at 37 °C, 5% CO_2_ and 95% humidity. The next day, the cells were washed with fresh medium and maintained until the next passage.

### Intestinal organoid differentiation

Human Intestinal organoids (IOs) were differentiated from hiPSCs using an adapted published protocol (32,62). Briefly, iPSCs were induced to form 3D IOs with a three-step protocol. On day 0, human iPSCs were induced to undergo definitive endoderm differentiation using RMPI1640 (Gibco) supplemented with Activin A (100 ng/ml, RnD). The next day, the medium was changed and fresh RMPI1640 supplemented with Activin A (100 ng/ml) and 0.2% HyClone-defined fetal bovine serum (FBS, GE Healthcare Bio-Sciences) was added. On the 3^rd^ and 4^th^ days, the medium was changed and RMPI1640 supplemented with Activin A (100 ng/ml) and 2% HyClone-defined FBS was added. The definitive endoderm was induced to mid- and hindgut differentiation by changing the medium daily [RPMI1640 supplemented with 15 mM HEPES, 2% HyClone-defined FBS, 500 ng/ml FGF4 (R&D), 500 ng/ml WNT3a (R&D)] until 3D spheroids were formed. Spheroids were collected, embedded in a drop of Cultrex Membrane Extract Type 2 (R&D), and fed with IO complete medium [Advanced DMEM F12 (Gibco) supplemented with B27 supplement (Thermo Fisher Scientific), GlutaMAX supplement (Thermo Fisher Scientific), penicillin and streptomycin (500 U/ml), 15 mM HEPES, 500 ng/ml R-spondin A (R&D), 100 ng/ml Noggin (R&D), 100 ng/ml EGF (R&D)], and the medium was changed twice a week. After approximately 50 days, the IOs were used in the experiments.

### Immunofluorescence staining

IOs were fixed with 4% paraformaldehyde (PFA) for 20 minutes at room temperature (RT) and washed three times with PBS. For whole-mount staining, the IOs were permeabilized according to the following protocol. For histological slides, IOs were first dehydrated using 15% sucrose overnight at 4 °C. Next, the samples were frozen in Tissue Freezing Medium (Leica Biosystems) in isopropanol cooled to −80°C and stored in freezer at −80°C. Next, the frozen samples were cut and rehydrated in PBS for 10 minutes. IO sections were permeabilized with PBS + 0.5% Triton-X100 for 15 minutes at RT and washed three times with IF buffer (PBS + 0.2% Triton-X100 + 0.05% Tween-20). Samples were blocked with IF buffer + 2.5% bovine serum albumin (BSA) for 1 hour at RT. The cells were labeled with primary antibodies (dilutions and clones are listed in Supplementary Table 1) overnight at 4 °C, washed three times with IF buffer, and incubated with secondary antibodies and phalloidin in IF buffer with 1% BSA for 2 hours at RT. IOs were washed three times with IF buffer and stained with DAPI for 5 minutes at RT. Before imaging, IOs were washed with IF buffer. IO sections were imaged with a Zeiss LSM 780 confocal microscope using 10x magnification. The images were processed using ImageJ software.

### Immunohistochemistry

IOs were washed with PBS, fixed with 4% PFA for 20 minutes at RT, and washed three times with PBS. The IOs were dehydrated using 15% sucrose overnight at 4 °C and frozen in Tissue Freezing Medium. The frozen samples were sectioned, and the sections rehydrated in PBS for 10 minutes. Endogenous peroxidases were blocked in 5% H_2_O_2_/methanol for 30 minutes at RT. Samples were washed in H_2_O and PBS and blocked with 1% BSA for 45 minutes at RT. The cells were labeled with primary antibodies (dilutions and clones are in Supplementary Table 1) overnight at 4 °C, washed three times with PBS, and incubated with biotinylated secondary antibodies (Supplementary Table 1) for 60 minutes at RT. The samples were washed, and the Vectastain® ABC-HRP Kit (Vector Laboratories) ImmPACT® DAB Substrate Kit (Vector Laboratories) were used to develop the signal. Next, the samples were counterstained with hematoxylin solution for 5 minutes at RT, then dehydrated using ethanol and mounted in Eukitt® Quick-hardening mounting medium (Sigma-Aldrich). Samples were imaged using a Zeiss Axio Scan Z1 slide scanner.

### IO dissociation and flow cytometry analysis

IOs were washed with cold Hank’s balanced salt solution (HBSS) to remove the Cultrex and cut with scissors into small pieces. The cells were dissociated with TrypLE (Gibco) for 10 minutes at 37 °C using constant shaking. Every 5 minutes, the samples were pipetted to dissociate the cells completely. The suspension of cells was filtered using a 70-μm strainer and washed with cold HBSS + 2% FBS. The cells were spun at 300 × *g* for 10 minutes at 4 °C and resuspended in staining buffer (PBS + 0.5% FBS + 2mM EDTA). The cells were labelled with fluorochrome-conjugated antibodies for 30 minutes on ice (dilutions and clones in Supplementary Table 1), washed with MACs buffer, and injected into a spectral flow cytometer SONY SA3800 (Sony Biotechnology). Data were analyzed using FlowJo v11 software (BD Life Sciences).

### Stimulation of IOs

IOs older than 50 days were stimulated with IO medium containing chemical triggers but without growth factors. To study the effect of neuromodulators on intestinal tissue, serotonin hydrochloride 10 µM (R&D), dopamine hydrochloride 10 µM (R&D), noradrenaline bitartrate 10 µM (R&D), and acetylcholine chloride 10 µM (R&D) were used. TNFα (10 ng/ml, R&D) was used to induce a pro-inflammatory milieu. For analysis using qPCR, and RNAseq IOs were stimulated for 4 hours. For fluorescent staining and chemotaxis experiments, the organoids were stimulated for 24 hours. The untreated control was cultivated with fresh IO medium without triggers for the appropriate time.

### RNA extraction and gene expression

Cells were lysed in TRI Reagent® (Merck), and RNA was isolated using RNaeasy Plus Micro kit (Qiagen) according to the manufacturer’s instructions. The isolated RNA was used for RNAseq, RT2 Profiler PCR Array, or qPCR. Prior the qPCR analysis, RNA was transcribed to cDNA using High-Capacity cDNA Reverse Transcription Kit (Applied Biosystems™) according to manufacturer’s protocol. qPCR analysis was done with TaqMan^TM^ TLR2, TLR3, TLR4, TLR5, TLR6, TLR7, TLR8, and TLR9 probes (details in Supplementary Table 2) and TaqMan^TM^ Gene expression Master Mix (Thermo Fisher Scientific) according to the manufacturer’s protocol. StepOne^TM^ Real-Time PCR System (Applied Biosystems) was used for qPCR analysis.

### RT^2^ Profiler PCR Array

RNA was isolated from IOs according to the aforementioned protocol. The cDNA was synthesized using RT^2^ First Strand Kit (Qiagen). To screen the expression profiles, RT^2^ Profiler PCR Array Human Inflammatory Response & Autoimmunity (Qiagen) was used according to the manufacturer’s protocol. The LightCycler® 480 Instrument II (Roche) was used for qPCR analysis.

### RNA sequencing

The integrity of the isolated RNA (according to the aforementioned protocol) was measured with a Fragment Analyzer using RNA Kit 15 nt (Agilent Technologies). Samples with an RNA integrity number ≥ 8 were used for sequencing. Then, 120 ng of total RNA was used as input for library preparation using QuantSeq 3ʹ mRNA-Seq FWD with UDI 12 nt Kit (v.2) (Lexogen) in combination with UMI Second Strand Synthesis Module for QuantSeq FWD. Quality control for library quantity and size distribution was done using the QuantiFluor dsDNA System (Promega) and High Sensitivity NGS Fragment Analysis Kit (Agilent Technologies). The final library pool was sequenced on an NovaSeq 6000 (Illumina) using the S4 Reagent Kit v1.5 300 with cycles in pair-end mode, resulting in an average of 20 million reads per sample.

### RNAseq analysis

Raw reads were quality-checked (FastQC) and preprocessed for adaptor trimming (Trim Galore). The trimmed reads were then mapped (HISAT2, Samtools) to the human genome (genome version: Ensembl GRCh38), demonstrating a total alignment rate of 45%-55%. Only uniquely mapped reads were retained for downstream processing. The strandedness and counts of each gene were determined (HTseq). All downstream procedures were performed in the R v4.4.3 environment. Differential expression analysis was performed with the DESeq2 pipeline. The differential expression results were filtered for low counts, and genes were considered differentially expressed when they demonstrated a |log2(FoldChange)| ≥ 0.6 and p-value ≤ 0.05. Gene ontology (GO) and gene set enrichment analysis (GSEA) were performed with the package clusterProfiler and the desktop version of GSEA MSigDB, respectively. Visualization of the results was performed in the R v4.4.3 environment (ggplot2, Complexheatmap).

### Peripheral blood mononuclear cell isolation

Peripheral blood mononuclear cells (PBMCs) were isolated from fresh healthy buffy coats from human blood samples (Department of Transfusion & Tissue Medicine of the Brno University Hospital). Human whole-blood samples were diluted with 2% FBS in PBS prior to gradient centrifugation on Lymphoprep (StemCell Technologies). A percentage of the freshly isolated PBMCs was labelled with anti-human CD3, CD4, CD8, CD14, CD16, CD19, and CD45 antibodies (details in Supplementary Table 1), referred to as “baseline” in the results. The rest of the PMBCs were used for chemotaxis/migration assays.

### Monocyte isolation

Monocytes were isolated from healthy human donor buffy coat (Department of Transfusion & Tissue Medicine of the Brno University Hospital) using RosetteSep^TM^ Human Monocyte Enrichment Cocktail (StemCell Technologies) according to the manufacturer’s protocol. Monocyte purity was assessed by flow cytometry using CD14, CD16, CD45, and HLA-DR markers (details in Supplementary Table 1), and dead cells were excluded using LIVE/DEAD™ Fixable Green Dead Cell Stain Kit (Invitrogen).

### Chemotaxis/migration assay

PBMCs isolated using the aforementioned protocol were used for chemotaxis/migration assays. The cells were seeded into the upper compartments of Transwell inserts (cellQUART, 5 µm pores) filled with Advanced DMEM F12 medium supplemented with B27 supplement, GlutaMAX supplement, penicillin and streptomycin (500 U/ml), and 15 mM HEPES. The lower compartments were filled with supernatants from IOs treated with 10 ng/ml TNFα, 10 µM serotonin hydrochloride, or a combination of both, for 24 hours. After 2 hours of incubation, the cells that had migrated to the lower compartment were collected, washed, and labelled with anti-human CD3, CD4, CD8, CD14, CD16, CD19, and CD45 antibodies (details in Supplementary Table 1). Dead cells were excluded using the LIVE/DEAD™ Fixable Violet Dead Cell Stain Kit (Invitrogen). The cells were analyzed using a spectral flow cytometer SONY SA3800 (Sony Biotechnology).

### Cocultivation of IOs and monocytes

Firstly, IOs were stimulated with 10 ng/ml TNFα, 10 µM serotonin hydrochloride, or a combination of both, for 24 hours, then extensively washed with cold PBS. Monocytes isolated using the aforementioned protocol were mixed with the IOs (10^5^ monocytes per organoid) by seeding in a drop of Cultrex. After 6 hours, organoids were extensively washed with cold PBS and dissociated according to the aforementioned protocol. The single cells in suspension were labelled with anti-human CD14, CD16, CD36, CD45, CD86, and HLA-DR antibodies (details in Supplementary Table 1). Dead cells were excluded using 7-AAD Viability Staining Solution (eBioscience). The cells were analyzed using a spectral flow cytometer SONY SA3800. Data were analyzed using FlowJo v11 software. For IF staining, monocytes and IOs were co-cultivated for 24 hours, and after extensive washing, fixed with 4% PFA for 20 minutes at RT. The fixed samples were used for histological slide preparation according to the protocol described above. The samples were stained with anti-human CD45, CD90, and E-cadherin antibodies (details in Supplementary Table 1).

### Statistical analyses

Statistical analyses were performed with GraphPad Prism 9, unless otherwise indicated. The results are shown as mean and SEM. Data were tested for normality before parametric or nonparametric statistical tests were applied.

