## Supplementary information for "Serotonin attenuates tumor-necrosis-factor-alpha-induced intestinal inflammation by interacting with human mucosal tissue"

This Supplementary information contains Supplementary Table 1 and Table 2, showing details of used antibodies and TaqMan^TM^ probes.

**Supplementary table 1. Details of used antibodies**

| **Antibody** | **Conjugate** | **Clone** | **Dilution** | **Vendor** | **Host** |
| --- | --- | --- | --- | --- | --- |
| **CD11b** | BV510 | ICRF44 | 1:100 | BioLegend | Mouse |
| **CD14** | PE | 63D3 | 1:100 | BioLegend | Mouse |
| **CD16** | BV711 | 3G8 | 1:100 | SONY | Mouse |
| **CD16** | eFluor 450 | eBioCB16 | 1:100 | eBioscience | Mouse |
| **CD19** | Biotin | HUB19 | 1:100 | eBioscience | Mouse |
| **CD3** | BV650 | UCHT1 | 1:100 | BD Horizon | Mouse |
| **CD36** | PE-Cy7 | 5-271 | 1:100 | BioLegend | Mouse |
| **CD4** | BV510 | SK3 | 1:100 | BioLegend | Mouse |
| **CD45** | BV421 | 2D1 | 1:100 | BioLegend | Mouse |
| **CD45** | PE-Cy7 | 2D1 | 1:100 | BioLegend | Mouse |
| **CD8** | BV421 | RPA-T8 | 1:100 | BioLegend | Mouse |
| **CD86** | BV711 | IT2.2 | 1:100 | BioLegend | Mouse |
| **CD90** | - | Polyclonal | 1:100 | R&D | Sheep |
| **CD90** | PerCP-Cy5.5 | eBio5E10 | 1:100 | eBioscience | Mouse |
| **CGA** | - | LK2H10 | 1:100 | Invitrogen | Mouse |
| **E-cadherin** | - | 24E10 | 1:300 | Cell Signaling TECHNOLOGY | Rabbit |
| **E-cadherin** | Alexa Fluor 488 | DECMA-1 | 1:300 | eBioscience | Rat |
| **EpCam(CD326)** | Biotin | 1B7 | 1:200 | eBioscience | Mouse |
| **HLA-DR** | PE-Dazzle 594 | L243 | 1:100 | BioLegend | Mouse |
| **Lysozym** | - | SB1 | 1:100 | Invitrogen | Mouse |
| **MUC5AC** | - | 45M1 | 1:500 | Abcam | Mouse |
| **Secondary anti mouse** | Alexa Fluor 546 | Polyclonal | 1:500 | Invitrogen | Goat |
| **Secondary anti mouse** | Alexa Fluor 647 | Polyclonal | 1:500 | Invitrogen | Donkey |
| **Secondary anti mouse** | Biotin | Polyclonal | 1:500 | Jackson Immuno Research | Goat |
| **Secondary anti rabbit** | Alexa Fluor 488 | Polyclonal | 1:500 | Invitrogen | Donkey |
| **Secondary anti rabbit** | Alexa Fluor 555 | Polyclonal | 1:500 | Invitrogen | Donkey |
| **Secondary anti Rat** | Alexa Fluor 488 | Polyclonal | 1:500 | Invitrogen | Donkey |
| **Secondary anti sheep** | Alexa Fluor 546 | Polyclonal | 1:500 | Invitrogen | Donkey |
| **Streptavidin** | Alexa Fluor 514 | - | 1:500 | Invitrogen | - |
| **Streptavidin** | APC | - | 1:500 | BioLegend | - |
| **TLR2** | - | D7G9Z | 1:200 | Cell Signaling TECHNOLOGY | Rabbit |
| **TLR3** | PE | TLR3.7 | 1:200 | eBioscience | Mouse |
| **TLR4** | - | 76B357.1 | 1:200 | Invitrogen | Mouse |
| **TLR6** | - | Polyclonal | 1:100 | Sigma | Rabbit |

**Supplementary table 2. Details of used TaqMan^TM^ probes**

| **Gene ID** | **TaqMan Probe ID** |
| --- | --- |
| **TLR2** | Hs00610101_m1 |
| **TLR3** | Hs01551078_m1 |
| **TLR4** | Hs00152939_m1 |
| **TLR5** | Hs01920773_s1 |
| **TLR6** | Hs01039989_s1 |
| **TLR7** | Hs00152971_m1 |
| **TLR8** | Hs07292888_s1 |
| **TLR9** | Hs00370913_s1 |
